# locStra: Fast analysis of regional/global stratification in whole genome sequencing (WGS) studies

**DOI:** 10.1101/2020.03.06.981050

**Authors:** Georg Hahn, Sharon M. Lutz, Julian Hecker, Dmitry Prokopenko, Michael H. Cho, Edwin K. Silverman, Scott T. Weiss, Christoph Lange, the NHLBI Trans-Omics for Precision Medicine (TOPMed) Consortium

## Abstract

*locStra* is an *R*-package for the analysis of regional and global population stratification in whole genome sequencing studies, where regional stratification refers to the substructure defined by the loci in a particular region on the genome. Population substructure can be assessed based on the genetic covariance matrix, the genomic relationship matrix, and the unweighted/weighted genetic Jaccard similarity matrix. Using a sliding window approach, the regional similarity matrices are compared to the global ones, based on user-defined window sizes and metrics, e.g. the correlation between regional and global eigenvectors. An algorithm for the specification of the window size is provided. As the implementation fully exploits sparse matrix algebra and is written in C++, the analysis is highly efficient. Even on single cores, for realistic study sizes (several thousand subjects, several million RVs per subject), the runtime for the genome-wide computation of all regional similarity matrices does typically not exceed one hour, enabling an unprecedented investigation of regional stratification across the entire genome. The package is applied to three WGS studies, illustrating the varying patterns of regional substructure across the genome and its beneficial effects on association testing.

## 1 Introduction

Genetic association studies are a popular mapping tool; however, they can be vulnerable to confounding due to population substructure (Laird and Lange, 2010). Numerous methods have been proposed to address this issue (Devlin and Roeder, 1999; Pritchard et al., 2000). Popular approaches rely on the genetic covariance matrix of the genotype data: EIGENSTRAT, STRATSCORE, multi-dimensional scaling (Price et al., 2006; Patterson et al., 2006; Lee et al., 2012), or on the genomic relationship matrix (Yang et al., 2011). For populations with recent admixture where each subject contains different proportions of the ancestral genomes, local ancestry-approaches have been suggested, e.g. RFMix (Maples et al., 2013), LAMP (Sankararaman et al., 2008), HAPMIX (Price et al., 2009), SABER (Tang et al., 2006), or ADMIXMAP (McKeigue et al., 2000). While such software packages are able to estimate local ancestry, they require the availability of the reference populations. Approaches as the ones developed by Wang et al. (2011) can incorporate the local ancestry information into the association testing framework.

While there is strong evidence for regional stratification (Martin et al., 2018; Zhong et al., 2019), the aforementioned matrix-based approaches are typically computed only globally. For the validity of the matrix-based approaches, it is only required that the selected loci are not in linkage disequilibrium (LD) (Laird and Lange, 2010). There are no theoretical constraints as to whether the loci are selected genome-wide or from a specific region. However, the computational burden has generally been prohibitive to use existing implementations for a genome-wide analysis of regional stratification. As most matrix-based approaches are designed for common, uncorrelated variant data, i.e. loci that are not in LD, many genomic regions do not contain a sufficient number of such loci for a regional computation of matrix-based approaches.

With the arrival of whole genome sequencing data, an abundance of data on densely spaced rare variants (RVs) that are mostly not in LD became generally available. As RVs are often population specific, they can be highly informative about population substructure and recent admixture (Bodmer and Bonilla, 2008; Kryukov et al., 2009; Keinan and Clark, 2012). Consequently, approaches based on Jaccard similarity matrices that utilize RV/WGS data have been developed (Prokopenko et al., 2016; Schlauch et al., 2017). However, the computational bottleneck has persisted for these approaches as well.

We developed *locStra*, an *R*-package implementing four approaches to assess population stratification in RVs at the regional and global level using (1) the genetic covariance matrix, (2) the genomic relationship matrix, (3) the unweighted and (4) weighted Jaccard similarity matrices (the weighted Jaccard matrix is also sometimes called s-matrix). The package is entirely written in C++, and all similarity matrices are algebraically transformed so that the computations are executed on sparse data structures. The sparse matrix structure is maintained throughout all computations to maximize computational efficiency. As a result, the runtimes for computing genetic similarity matrices are substantially shorter for *locStra* than for existing packages. Using sliding windows (Morrison et al., 2013; Panoutsopoulou et al., 2013; Yazdani et al., 2015) of user-specified length, *locStra* enables the fast analysis of regional stratification at the genome-wide level, even on desktop computers.

The article is structured as follows. Section 2 briefly introduces the methodology we employ in this work, in particular we describe the functionality of our software. Results are presented in Section 3, investigating amongst others the regional stratification in data from the 1,000 Genome Project (Section 3.1), a comparison to PLINK2 (Section 3.1.3), and the selection of suitable window sizes (Section 3.1.4). Moreover, we visualize regional substructures using a population isolate (Section 3.2) and demonstrate the beneficial effects of regional stratification on association testing by correcting a linear regression using global and regional principal components (Section 3.3). The article concludes with a discussion in Section 4. The appendix highlights certain details of our optimized implementations including theoretical runtimes (Section A).

## 2 Methods

### 2.1 Software description

The core implementation of *locStra* is based on fully sparse matrix algebra in C++, using *RcppEigen* of Bates and Eddelbuettel (2013). The package is available on *The Comprehensive R Archive Network*, see Hahn et al. (2020).

The *locStra* package makes a total of seven functions available which are described in the following subsections.

#### 2.1.1 Dense and sparse matrix implementations

Four functions provide C++ implementations of standard approaches to population stratification, both for dense and sparse matrix algebra. The code handles dense and sparse input matrices separately since either version can be inefficient if used for matrices of the wrong type. Each of the functions listed below takes at least two arguments: the input matrix (which can be in dense or sparse format) on which to compute the similarity measure, and a boolean argument *dense* to switch between C++ implementations for dense and sparse matrix algebra. The default is *dense*=*False*. The input matrix always contains the genotype/dosage data oriented as number of variants/rows by number of people/columns.

1. The function *covMatrix* computes the genetic covariance matrix (Price et al., 2006). The entries of the input are allowed to be any real valued numbers.
2. The function *grMatrix* computes the genomic relationship matrix (GRM) as defined in Yang et al. (2011). The input must be a binary matrix. Both the classic and robust versions (Wang et al., 2017) of the GRM are supported, and can be switched using the boolean flag *robust*. The default is *robust*=*True*.
3. The function *jaccardMatrix* computes the Jaccard similarity matrix (Prokopenko et al., 2016). The input must be a binary matrix, where a one indicates the presence of a variant.
4. The function *sMatrix* implements the weighted Jaccard matrix (Schlauch et al., 2017; Schlauch, 2016). In addition to the boolean *dense* argument, the function *sMatrix* also has a boolean argument *phased* to indicate if the input data are phased (default is *phased*=*False*). The input can be any real valued matrix. The last argument is the integer *minVariants* which is a cutoff value for the minimal number of variants to consider (default is *minVariants*=*5*). If no variants remain after applying the *minVariants* cutoff, the output will be a matrix with *NA*.

The unweighted and weighted Jaccard indices, traditionally a similarity index for sets, are two recently proposed approaches for the analysis of rare variant data which were shown to provide a higher resolution than the other approaches (Prokopenko et al., 2016). The entries of the Jaccard matrix measure the set-theoretic similarity of the genomic data of all pairs of subjects, and can be computed efficiently using only binary operations.

#### 2.1.2 Main function

The main function of the package is *fullscan*. It has five arguments and allows for a flexible specification of the regional population stratification scan of the data through its generic structure.

- The first input is the (sparse) matrix containing the sequencing data. The input matrix is assumed to be oriented to contain the data for each individual per column, thus the dimension of the input is the number of variants by the number of people.
- The second argument is a two-column matrix (called *windows*) that contains the window specification of the scan. The window matrix has as many rows as there are windows to iterate over, and two columns. The two columns are the start and end positions of each window. The matrix of sliding windows can easily be generated with the auxiliary function *makeWindows* described in the next subsection.
- The third argument, *matrixFunction*, handles the processing of each sliding window. The function takes one input argument (often a matrix). Any function can in principle be used. Typical choices are *covMatrix*, *grMatrix*, *jaccardMatrix*, or *sMatrix*.
- Next, the modular structure of *fullscan* requires the specification of a *summaryFunction* for the processed data before comparison. This can be any function of one input argument that is compatible with the output of *matrixFunction*. As an example, one might want to set the *matrixFunction* to *covMatrix*, which returns a square similarity matrix, and accordingly define the *summaryFunction* to compute the largest eigenvector on its square matrix input. This can easily be done with the function *powerMethod* described in the next subsection.
- *fullscan* uses its fifth input argument, the function *comparisonFunction*, to compare summaries (e.g., first eigenvectors) on a regional and a global level. The *comparisonFunction* must have two arguments as input, both of which need to be compatible with the output of the function *summaryFunction*. For instance, given the *summaryFunction* computes the first eigenvector, we might want to compare the first eigenvectors on a global and regional level by means of the native *R* correlation function *cor* for two vectors.

The output of *fullscan* is a two column matrix with global and regional comparison values per row, where each row corresponds to a row (and thus a window) in matrix *windows* in the same order.

#### 2.1.3 Auxiliary functions

Two functions provide additional functionality:

1. The function *makeWindows* generates a two-column matrix of non-overlapping or overlapping windows for the main function *fullscan*. The function takes as its arguments the length of the data, the window size and an offset. If the offset is set equal to the window size, non-overlapping windows are obtained. If the offset is less than the window size, sliding windows of given window size and offset are obtained.
2. The function *powerMethod* provides a C++ implementation of the power method for fast iterative computation of the largest eigenvector (von Mises and Pollaczek-Geiringer, 1929). The function can be used as *summaryFunction* in the main function *fullscan*.

#### 2.1.4 Other comparison measures

The modular structure of *locStra* allows to specify (1) the similarity measure on the genome (the *matrixFunction*; for instance, the Jaccard matrix); (2) the summary statistic for the similarity matrix as function *summaryFunction*; and (3) a comparison measure on either the similarity matrices or the summary statistic (function *comparisonFunction*). In this work we always summarize the four similiarity matrices with the first eigenvector (as *summaryFunction*) and compare the correlation between eigenvectors (as *comparisonFunction*). However, many more sensible choices exist which include:

1. The similarity matrices can be compared directly using, for instance, the *L_p,q_*, Frobenius, maximum or Schatten matrix norms as *comparisonFunction*. In this case the *summaryFunction* is the identity function.
2. Apart from eigenvectors, two similarity matrices can be summarized using other traditional tools such as their eigenvalues, or the condition number of the difference between both which, if large, indicates that the matrices are close in this specific sense.
3. Apart from the first eigenvector, the similarity matrices can be summarized with a linear combination of higher order eigenvectors to capture more principal components. Moreover, the eigenvectors can be weighted with their corresponding eigenvalues.
4. Apart from using vector correlation, eigenvectors and other vector-valued measures can be compared using vector norms, the angle between them, etc.

However, some measures might be more meaningful than others depending on the context of the comparison and application. We did experiment with different measures and found the correlation between the first eigenvectors to capture best the variability within each chromosome.

## 3 Results

### 3.1 Regional stratification analysis of the 1,000 Genome Project

To illustrate the practical relevance of regional substructure analysis and its feasibility at the genome-wide level, we apply *locStra* to all chromosomes of the 1,000 Genome Project (The 1000 Genomes Project Consortium, 2015) and take a closer look at the results for four chromosomes, precisely chromosomes 5, 7, 12, and 16. Importantly, we investigate runtimes across all chromosomes, and present an approach to select suitable window sizes for population stratification.

Before applying *locStra*, the raw data from the 1,000 Genome Project is prepared using PLINK2 (Purcell and Chang, 2019) with cutoff value 0.01 for option *--max-maf* to select rare variants. We applied LD pruning with parameters *--indep-pairwise 2000 10 0.01*. Analysis results are shown for the European super population alone (503 subjects, ca. 5 million RVs) of the 1,000 Genome Project. All timings presented in this and the following sections were measured on one Intel QuadCore i5-7200 CPU with 2.5 GHz and 8 GiB of RAM.

Using a sliding window approach of 128, 000 RVs (as suggested by our window selection algorithm), we used *locStra* to compute the correlations between the first eigenvector of all regional similarity matrices with the corresponding first eigenvector of the global similarity matrix.

#### 3.1.1 Data analysis results for certain chromosomes of the 1,000 Genome Project

The regional substructure analysis in Figure 1 reveals several notable features. Regardless of the type of similarity matrix used for the regional substructure analysis, there are only a few genomic regions for which the regional and global substructures are similar in terms of the first eigenvectors. Overall, there is substantial variability of the regional substructure across the genome, when measured via the similarity matrices. This observation also has implications for association mapping, as the association analysis is typically adjusted for the global eigenvectors to minimize potential genetic confounding. It will be subject of future research to investigate the best ways to incorporate regional substructure adjustments based on RVs in genetic association testing.

**Figure 1:**
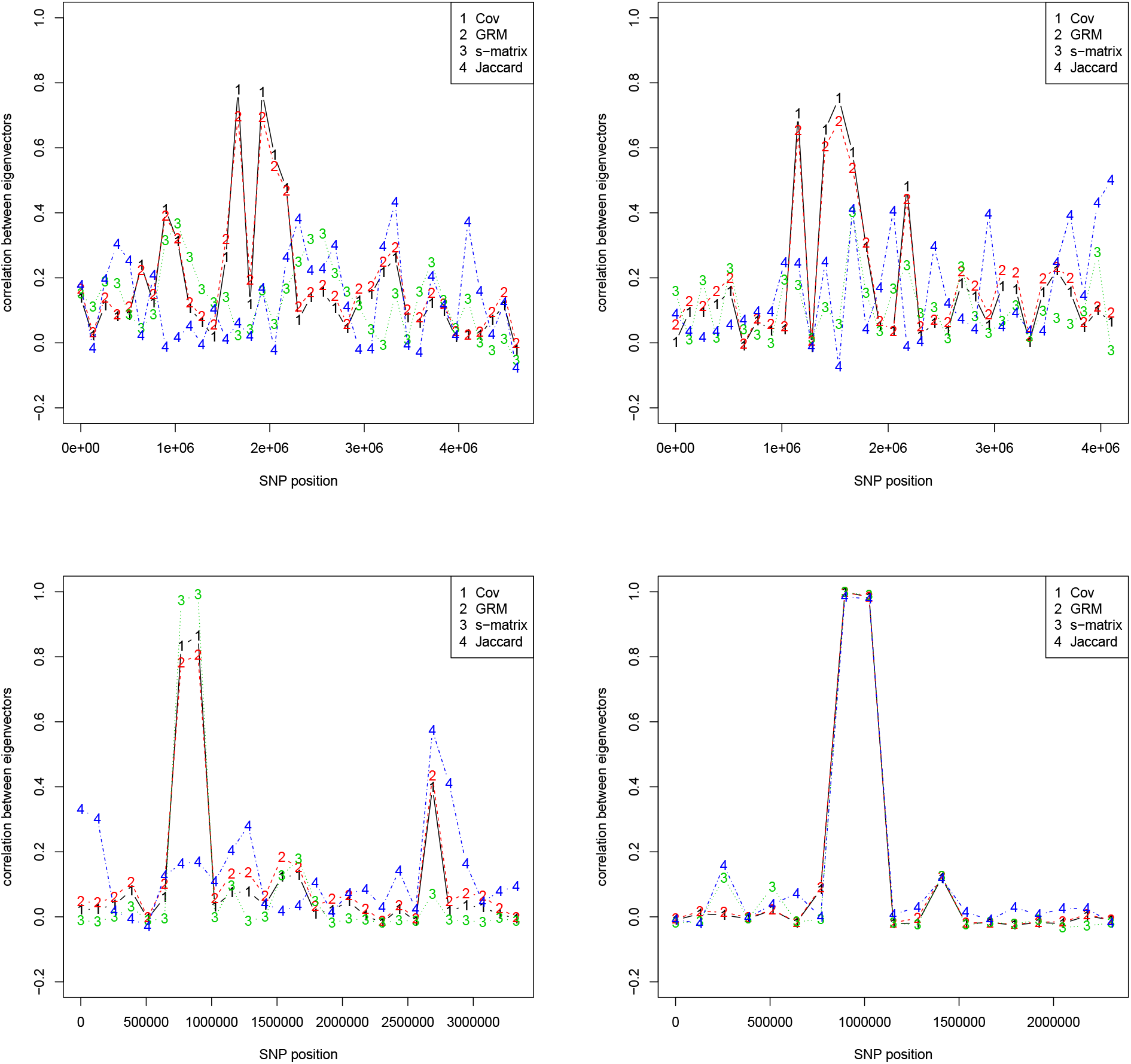
Rare variants for the super population EUR of the 1,000 Genomes Project. Correlation of regional to global eigenvectors for chromosomes 5 (top left), 7 (top right), 12 (bottom left), and 16 (bottom right). Covariance matrix, GRM matrix, s-matrix, and Jaccard matrix. Window size 128000 RVs.

In the areas where the correlation between the regional and global first eigenvector is not high, the standard Jaccard approach is able to maintain the highest correlation values compared to the other similarity matrices. In the areas where the regional first eigenvectors of Cov, GRM and s-matrix/weighted Jaccard are highly correlated with the corresponding global first eigenvectors, the first eigenvector of the standard Jaccard approach is often almost uncorrelated with the global one. Further methodological and substantive research is required to understand the reasons for these performance differences. It is part of our ongoing research and beyond the scope of this manuscript.

In order to contrast rare and common variants, Figure 2 shows similar plots for the same four chromosomes on common variants. We observe that, as expected, there are little to no variation across the genome.

**Figure 2:**
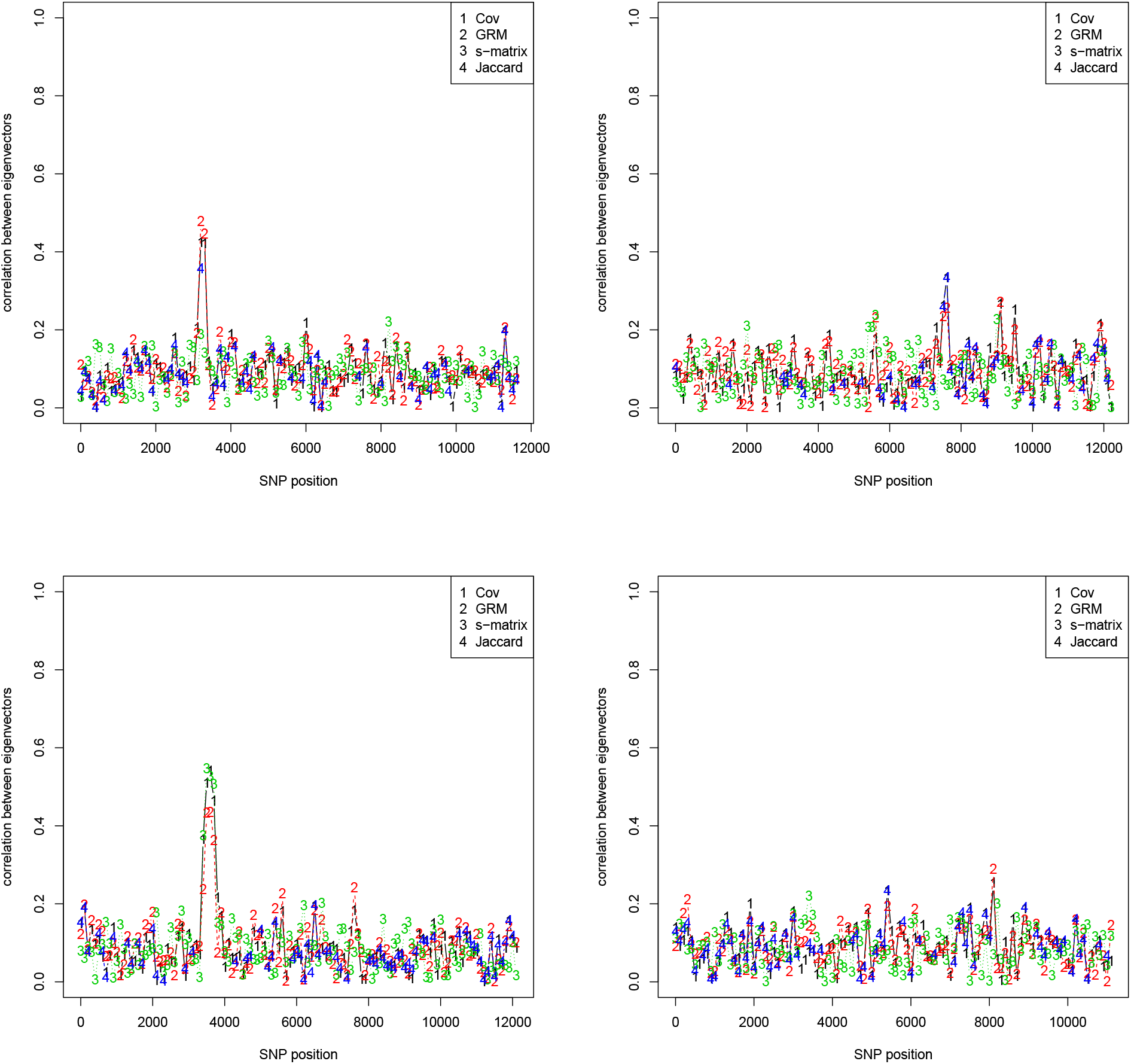
Common variants for the super population EUR of the 1,000 Genomes Project. Correlation of regional to global eigenvectors for chromosomes 5 (top left), 7 (top right), 12 (bottom left), and 16 (bottom right). Covariance matrix, GRM matrix, s-matrix, and Jaccard matrix. Window size 100 RVs.

#### 3.1.2 Runtime of *locStra* for the 1,000 Genome Project Analysis

Figure 3 shows the runtime in seconds for the *R* function *fullscan* (see Section 2.1.2) as a function of the window sizes. Each plot depicts the minimal and maximal runtime observed among any of the chromosomes per window size, as well as the mean runtime for a particular window size when averaged across all chromosomes.

**Figure 3:**
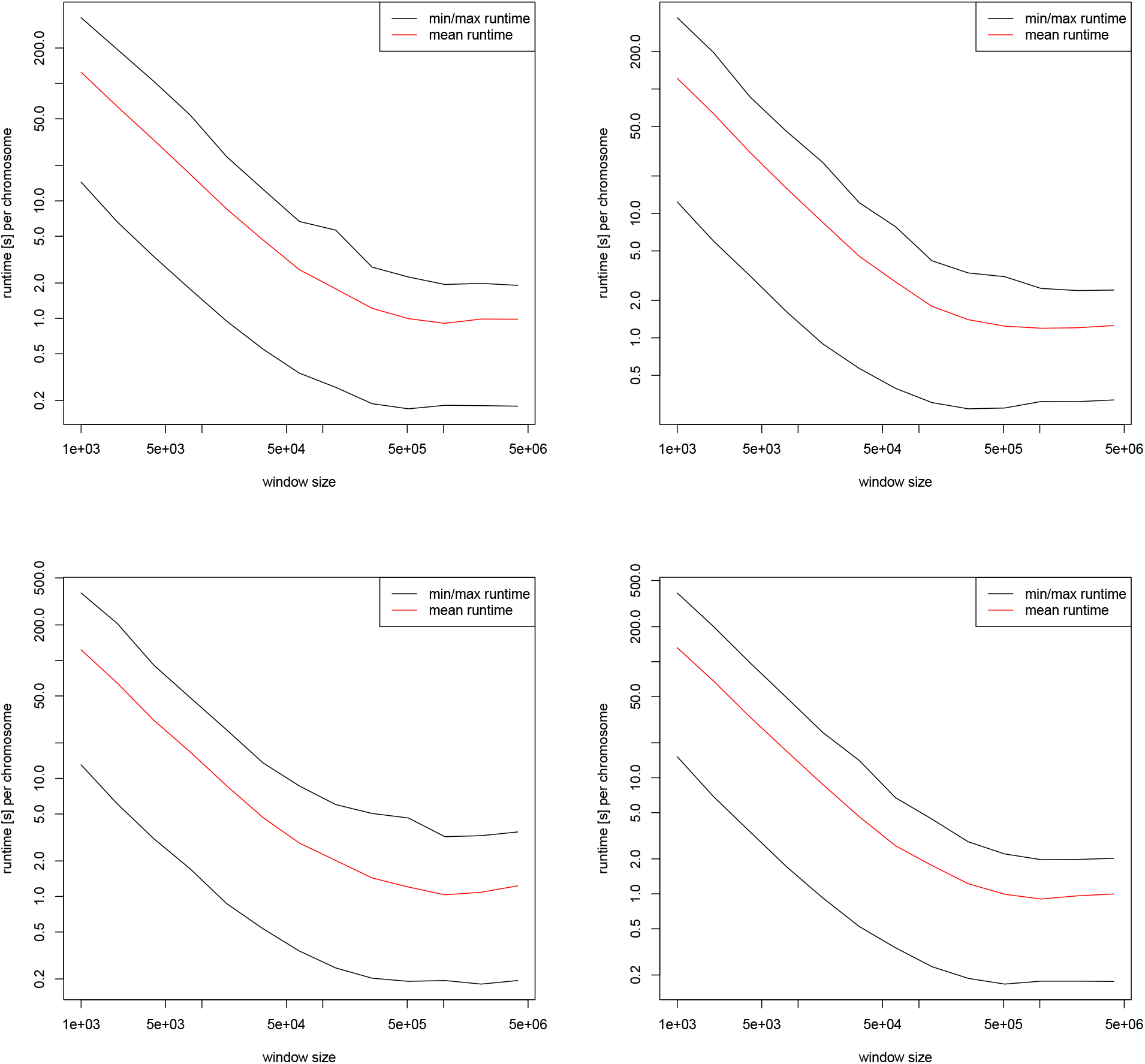
Super population EUR of the 1,000 Genomes Project. Runtime in seconds as a function of the window sizes across all chromosomes for the computation of the covariance matrix (top left), GRM matrix (top right), s-matrix (bottom left), and Jaccard matrix (bottom right). All plots show the minimal and maximal runtimes for any of the chromosomes, as well as the mean runtime averaged across all chromosomes. Logarithmic scale on the *x*- and *y*-axes.

One can see that the mean runtime never exceeds 500 seconds for a complete scan of any chromosome. As expected, the runtime decreases for larger window sizes. For a realistic window size of e.g. 10^4^ or 10^5^ (see Section 3.1.4), the runtime for any method is in the neighborhood of one minute for a full scan. Repeating the runtime analysis for the AFR super population group of the 1,000 Genome Project shows qualitative similar results.

#### 3.1.3 Comparison of *locStra* to PLINK2 on chromosome 1 of the 1,000 Genome Project

Performing a similar scan of the genome with PLINK2 (Purcell and Chang, 2019), which is one of the most frequently used tools to compute genetic variance/covariance matrices, is possible but has a considerably higher computational runtime. This is due to the fact that, although PLINK2 can both extract genomic regions specified by rs numbers and compute eigenvectors, it does so via file operations which are very slow, especially if called multiple times as it is needed for a genome-wide scan.

To compare *locStra* to PLINK2, we first prepare the data of chromosome 1 using the same parameters as in the previous subsections. Since *locStra* and PLINK2 require different input files, we write the curated data for chromosome 1 once in the .*bed* format for PLINK2 to read, and once convert it into a sparse matrix of class *Matrix* in *R* (saved as *.Rdata* file).

In PLINK2, the first eigenvector can be computed on the .*bed* file input with the option *--pca 1*, and in order to do a regional scan, a variant range on the data can be specified with the parameters *--from* and *--to* followed by the rs numbers. After computing the first global eigenvector or one regional eigenvector, PLINK2 writes the vector data into a file *.eigenvec*, from which the eigen-vectors are read in order to compute correlations between them. In this way, a complete scan of correlations between global and regional eigenvectors can be carried out in PLINK2.

For *locStra*, we load the sparse matrix input data into *R* and employ the function *fullscan* from the *R* package to carry out a complete scan.

Results for three different window sizes are given in Table 2, showing both the runtimes (in seconds) for the computation of the single global eigenvector on the full data, as well as for a complete scan (which includes the computation of the global eigenvector before starting the scan). Since the times for PLINK2 necessarily include the read/write operations for file in- and outputs, we likewise report times for *locStra* that include the reading time of the input data for a fair comparison. As visible from the table, *locStra* is at least one order of magnitude faster than PLINK2 on the data of the 1,000 Genome Project. Moreover, the speed-up seems to be more pronounced for larger window sizes. Based on further experiments (not reported here), we expect those results to generalize for larger datasets.

**Table 1:** TOPMed Omics Support Table. Broad Genomics = Broad Institute Genomics Platform; Broad Metabolomics = Broad Institute and Beth Israel Metabolomics Platform; Keck MGC = Keck Molecular Genomics Core Facility; NWGC = Northwest Genomics Center.

| TOPMed<br>Accession # | TOPMed<br>Project | Parent Study<br>Short Name | TOPMed<br>Phase | Omics Center | Omics Support | Omics Type |
| --- | --- | --- | --- | --- | --- | --- |
| phs001726 | CRA_CAMP | CAMP | 3 | NWGC | HHSN268201600032I | WGS |
| phs001726 | CRA_CAMP | CAMP | 5 | Broad Metabolomics | HHSN268201600034I | Metabolomics |
| phs001726 | CRA_CAMP | CAMP | 5 | Keck MGC | HHSN268201600038I | Methylomics |
| phs000951 | COPD | COPDGene | 1 | NWGC | 3R01HL089856-08S1 | WGS |
| phs000951 | COPD | COPDGene | 2 | Broad Genomics | HHSN268201500014C | WGS |
| phs000951 | COPD | COPDGene | 2.5 | Broad Genomics | HHSN268201500014C | WGS |
| phs000951 | COPD | COPDGene | 4 | NWGC | HHSN268201600032I | RNASeq |
| phs000951 | COPD | COPDGene | 5 | NWGC | HHSN268201600032I | Methylomics |
| phs000988 | CRA_CAMP | CRA | 1 | NWGC | 3R37HL066289-13S1 | WGS |
| phs000988 | CRA_CAMP | CRA | 3 | NWGC | HHSN268201600032I | WGS |
| phs000988 | CRA_CAMP | CRA | 5 | Broad Metabolomics | HHSN268201600034I | Metabolomics |
| phs000988 | CRA_CAMP | CRA | 5 | Keck MGC | HHSN268201600038I | Methylomics |

**Table 2:** Comparison of *locStra* and PLINK2. Runtimes in seconds for the computation of the global eigenvector (global EV) and for a full stratification scan of chromosome 1 of the 1,000 Genome Project as a function of the window size.

| window size | *locStra* |  | PLINK2 |  |
| --- | --- | --- | --- | --- |
|  | global EV | full scan | global EV | full scan |
| 1000 | 1.4 | 332.1 | 65.3 | 6343.6 |
| 10000 | 1.5 | 31.3 | 61.2 | 731.7 |
| 100000 | 1.5 | 4.9 | 66.7 | 189.1 |

Figure 4 shows how the results of PLINK2 and *locStra* compare. For this we compute the GRM similarity matrix on the EUR super population of the 1,000 Genome Project with both programs. We then plot the first two principle components against each other using the same x-axis and y-axis scaling. We observe that throughout our experiments (results for other chromosomes not shown), the stratification is similar, in particular the point clouds are located in roughly the same spots. However, the projection of *locStra* is often more spread out, making it easier to differentiate between the population subgroups contained in the dataset.

**Figure 4:**
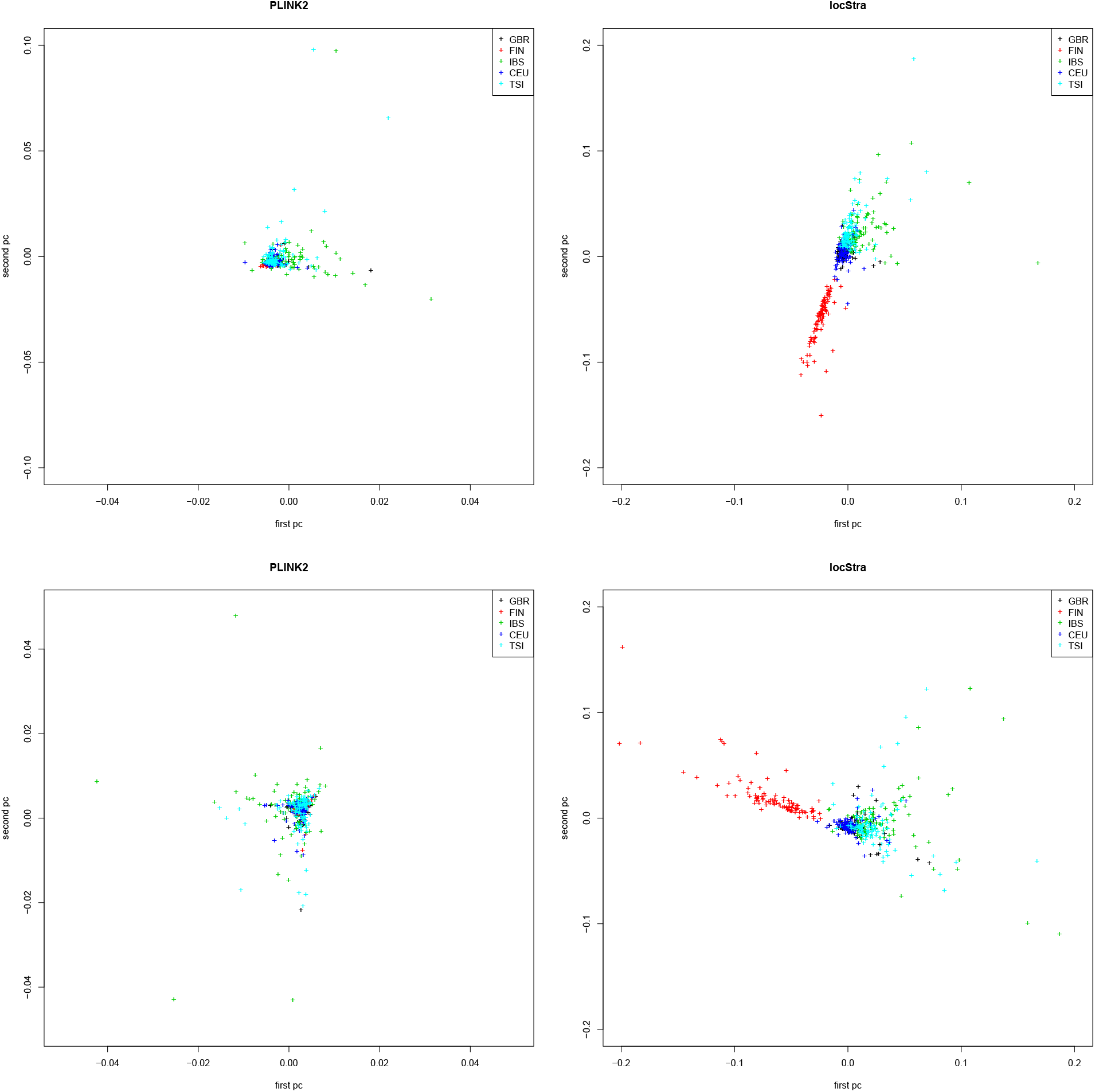
Super population EUR of the 1,000 Genomes Project with British (GBR), Finnish (FIN), Iberian (IBS), Utah resident (CEU) and Toscani (TSI) subgroups. First two principal components for the GRM similarity matrix computed with PLINK2 (left) and *locStra* (right) for chromosome 1 (top) and chromosome 6 (bottom).

#### 3.1.4 Selecting suitable window sizes for population stratification

An interesting question pertains to the selection of an appropriate window size for population stratification. Two quantities work against each other in the process of selecting a suitable window size: As the window size becomes larger, fewer windows are used in the scan of the data, and thus the correlation between regional and global eigenvectors increases as seen in Figure 5 (left). On the other hand, larger window sizes imply the usage of fewer windows in the scan, thus causing fewer data points to be calculated.

**Figure 5:**
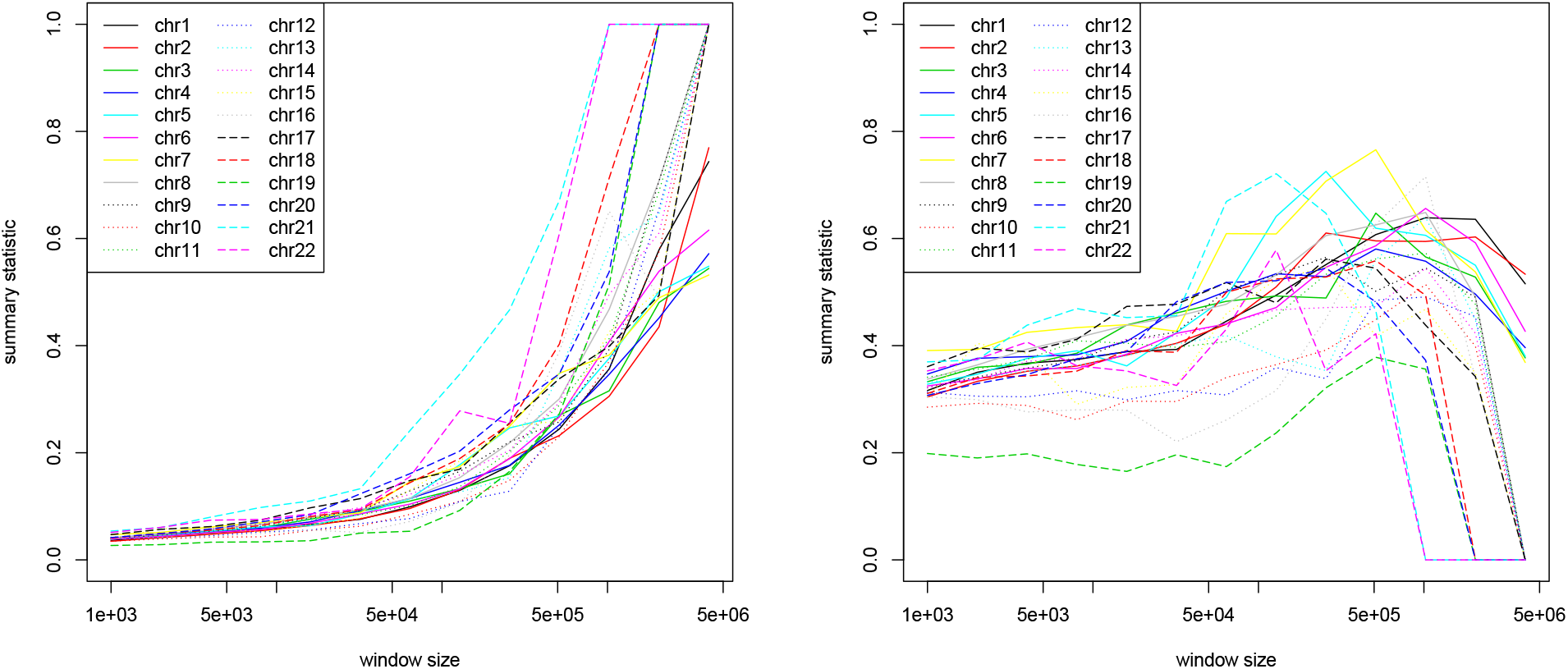
Super population EUR of the 1,000 Genomes Project. Left: Mean correlation across all windows as a function of the window size. Right: Mean correlation across all windows multiplied by the logarithm of the number of windows, again as a function of the window size. Input data are the correlations between global and regional eigenvectors of the Jaccard matrices for differrent window sizes.

This gives rise to a variety of heuristics which can be used to trade off the two quantities (correlation and number of data points), with the aim to arrive at a sensible (though not theoretically proven) measure indicating good selections for the window size. As an example, we multiply the mean correlation among all windows (for a particular window size) with the logarithm of the number of windows generated using that size (alternatively, any monotonic transformation of the number of windows can be used).

The resulting measure is displayed in Figure 5 (right) for the EUR super population, and in Figure 6 for the AFR super population of the 1,000 Genome Project. It can be seen that for very small and large window sizes, the heuristic measure is lower than in-between the two extremes. For both the EUR and AFR super populations of the 1,000 Genomes Project, we observe a peak at roughly a window size of 10^5^ RVs. Interestingly, the shape of the curves is almost identical across all chromosomes. This is attributed to the fact that the slopes in Figure 5 (left) and Figure 6 (left) is very similar for all chromosomes.

**Figure 6:**
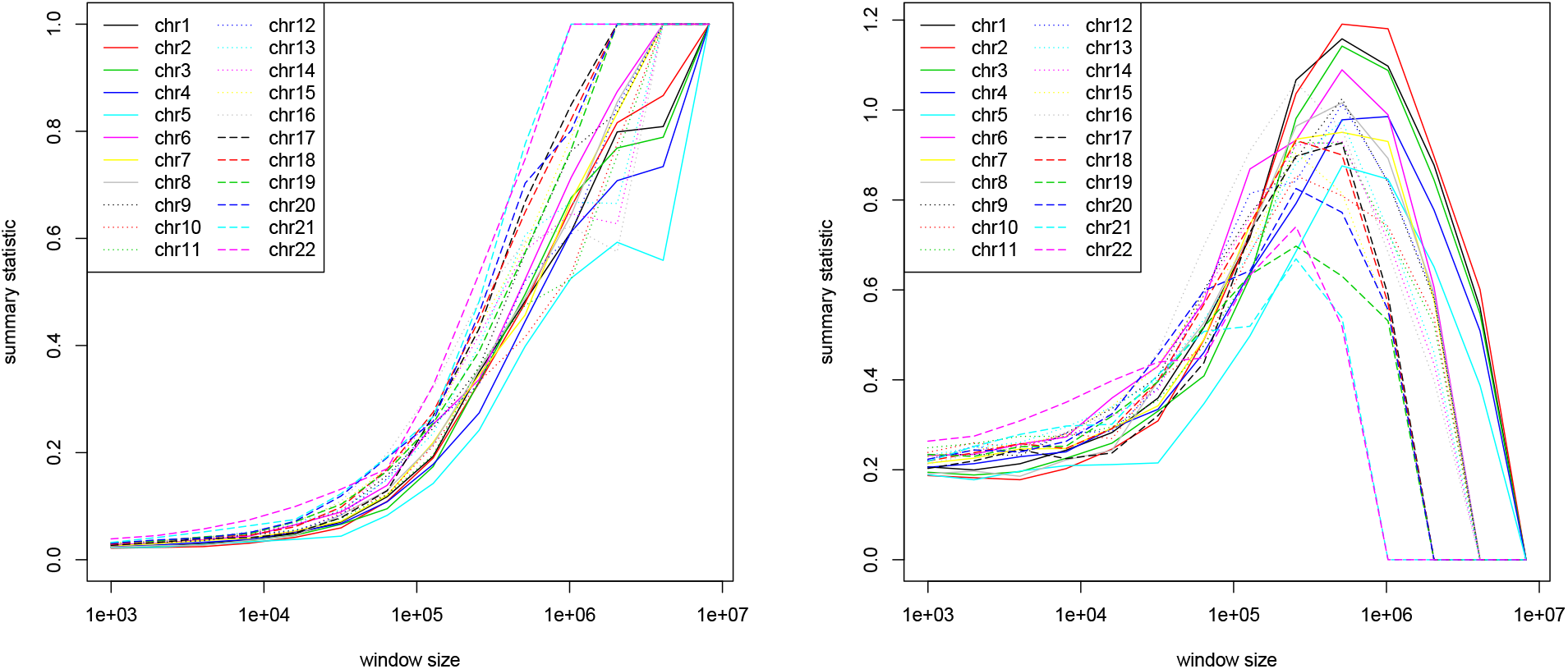
Super population AFR of the 1,000 Genomes Project. Setting as in Figure 5. Left: Mean correlation across all windows as a function of the window size. Right: Mean correlation across all windows multiplied by the logarithm of the number of windows, again as a function of the window size.

For the analysis of EUR and AFR super population data from the 1,000 Genomes Project, we will use a window size of around 10^5^ RVs. This idea of trading off the mean correlation with the number of data points generalizes to other datasets and can be seen as an heuristic guideline.

### 3.2 Data analysis of a Childhood Asthma Study from Costa Rica (population isolate)

We now analyze the global and the regional population substructure and the differences they exhibit on a cluster level in a dataset of a Costa Rica population isolate, as one expects stronger degrees of stratification in such a sample. The dataset includes children aged 6 to 14 and their parents (2736 subjects, 1824 of which are parents) from GACRS (Genetics of Asthma in Costa Rica Study) family-based trios recruited from a genetically homogeneous Hispanic population isolate living in the Central Valley of Costa Rica. This population has one of the highest prevalences of asthma in the world. Please see Hunninghake et al. (2007) for a detailed description of the recruitment process. The study has been sequenced as part of the TOPMED Project. The data are available through dbGaP (NHLBI TOPMed, 2019).

To avoid genetic correlations among study subjects due to family structure, we only select the parents for the analysis, and prepare their genetic data using PLINK2 with cutoff value 0.01 for option *--max-maf* to select rare variants. We applied LD pruning with parameters *--indep-pairwise 2000 10 0.01*.

To evaluate the population substructure in the data, we compute the Jaccard similarity matrix globally and regionally for one window (the middle window) on each chromosome. We selected here the Jaccard approach for ease of presentation. None of the qualitative conclusions that we reach below would have been different had we selected a different similarity matrix.

Figure 7 shows the first two “global” principal components of the Jaccard matrix, meaning the first two principal components of the Jaccard matrix which was computed globally based on subsampled loci from the entire genome. To be precise, to generate Figure 7, we randomly sampled 10^5^ RVs from each chromosome, and combined them into one matrix on which the Jaccard similarity measure was computed. The plot shows that most samples get mapped into a consistent point cloud. However, we observe that this cloud stretches out to the above and below, with three clear outliers marked in red.

**Figure 7:**
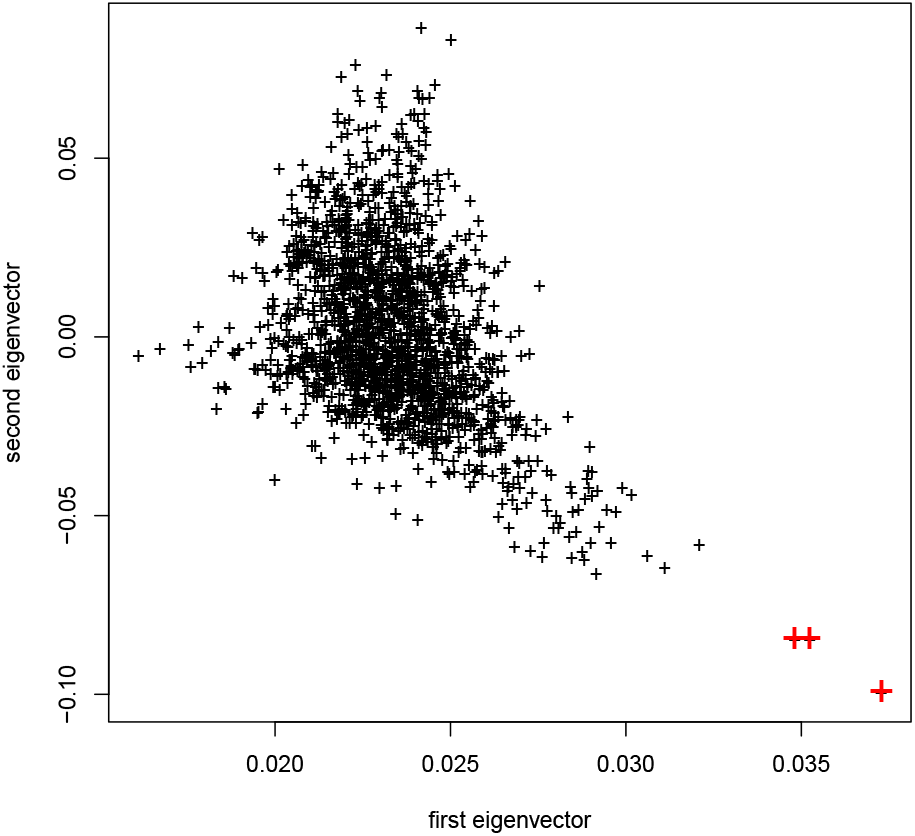
Costa Rica population isolate. First two principal components for the Jaccard similarity matrix. All chromosomes combined. Three outliers marked with red crosses.

For each of the 22 autosomal chromosomes, Figure 8 shows the plots of the first two “regional” principal components of the Jaccard matrix computed on the middle region (of size 10^5^ loci) on the chromosome. To generate each of the subplots in Figure 8, we selected a window size of 10^5^ for each chromosome (resulting in around 20 windows per chromosome depending on the size of the chromosome data), and computed the similarity matrix on the middle window.

**Figure 8:**
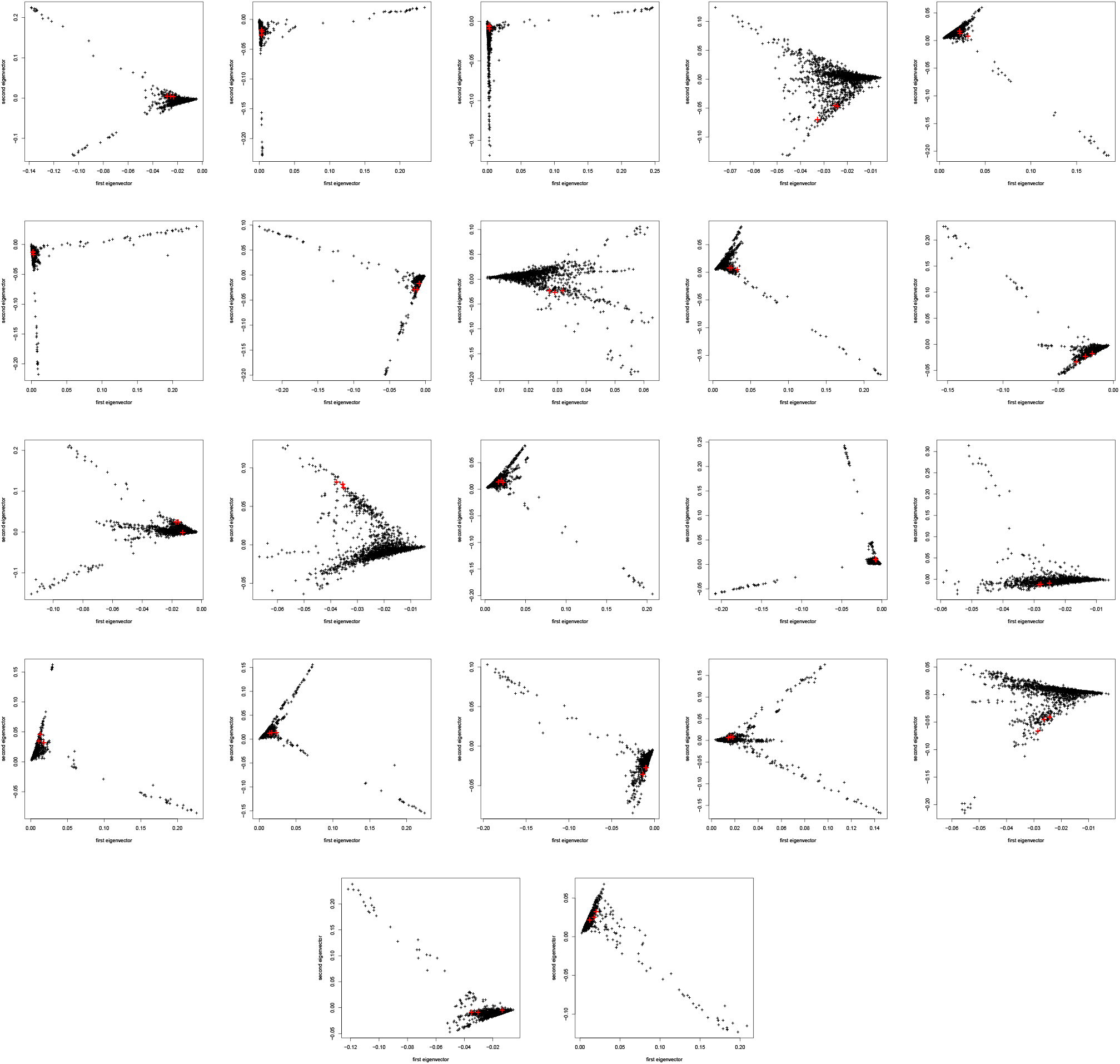
All 22 chromosomes of the Costa Rica population isolate. First two principal components for the Jaccard similarity matrix computed for the middle window of a stratification scan with window size 10^5^. Separate plot for each chromosome (starting with chromosome 1 in the top left corner and continuing in a row-wise fashion). The three outliers of Figure 7 are marked again in each subplot in red.

Interestingly, Figure 8 shows that on a regional level, the substructure can vary substantially. While it can be very similar to the global substructure for some of the regions, often it is more extreme or fundamentally different, showing sub-clusters that are not detectable in the global components. Moreover, we observe that on a regional level, the structure is often more extreme, exhibiting large branches that are substational proportion of samples fall into. By exemplarily marking the three outliers of Figure 7 in each of the 22 subplots (in red), we observe that each subplot exhibits are much finer structure than the global one, with outliers which are not detectable in a global analysis.

Analyzing the relationship of regional outliers among each other and with respect to the global PCA plot, and inferring possible conclusions from it, remains for further research. A first impression is given in Figure 9, which shows where the outliers from the branches observed in Figure 8 are located in the global PCA plot (all outliers from the 22 regional plots are displayed in Figure 9 with their chromosome number). We observe that there is indeed a structure: for instance, samples from the outliers of chromosomes 1, 8, and 15 seem to be located exclusively in the upper part of Figure 9.

**Figure 9:**
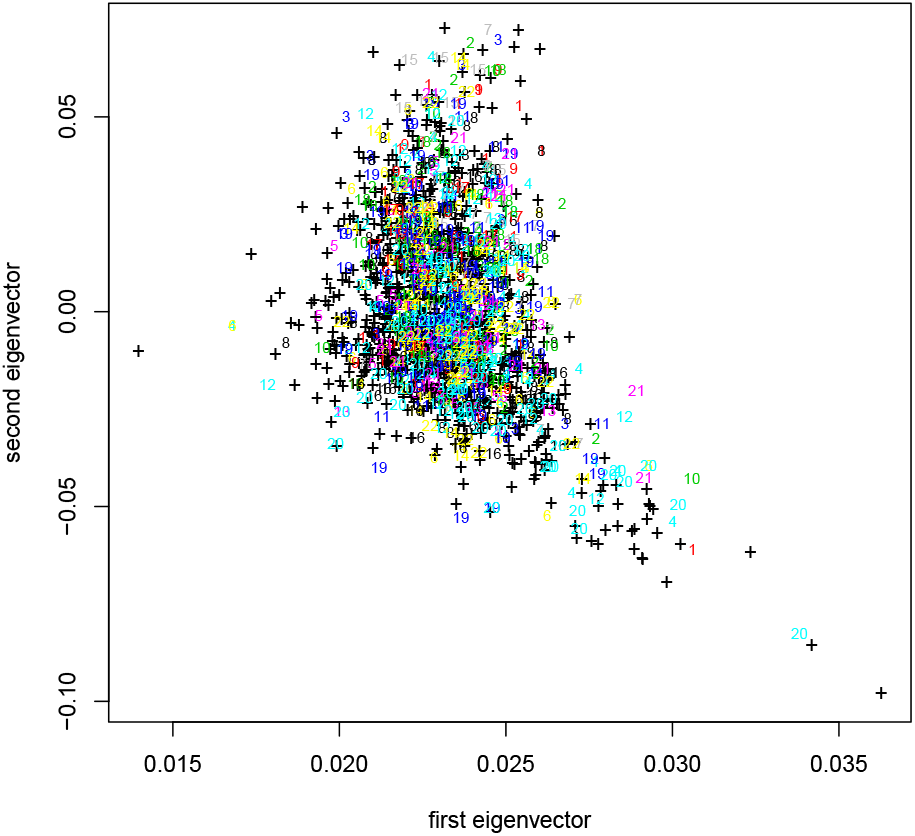
Costa Rica population isolate. First two principal components for the Jaccard similarity matrix. All chromosomes combined. The samples contained in the branches of the 22 regional plots of Figure 8 are both colored by chromosome, and labeled with their chromosome number.

The corresponding results for the other similarity matrix approaches support the same conclusion (data not shown). The findings of the Costa Rica data analysis clearly demonstrate the importance of regional substructure analysis and the utility of the proposed *locStra* package.

### 3.3 Correcting a linear regression with global and regional principal components

To assess the effects of regional substructure on association testing, we consider an example in the COPDGene study, a case-control study of Chronic Obstructive Pulmonary Disease (COPD) in current and former smokers (Regan E.A., 2010). The study has been sequenced as part of the TOPMED Project. The data are available through dbGaP (NHLBI TOPMed, 2018).

We examine the effect of the particular SNP *rs16969968* (chromosome 15) on FEV_1_ which is a well-established risk locus for COPD and cigarette smoking (Pillai et al., 2009; Lutz et al., 2015). It is unclear whether the regional substructure or the global substructure is more relevant for a particular locus that is tested for association. It is important to note that the inclusion of additional principal components will not have a major impact on the power of the association analysis, given the current sample size of such studies. As a consequence, the analysis plan is to evaluate three regression models:

- Model 1: Regress FEV_1_ on *rs16969968* adjusting for age, height, sex, and the first 5 global principal components;
- Model 2: Regress FEV_1_ on *rs16969968* adjusting for age, height, sex, and the first 5 regional principal components that are computed for the region that harbors *rs16969968*;
- Model 3: Regress FEV_1_ on *rs16969968* adjusting for age, height, sex, and the first 5 regional principal components and on the first 5 global principal components.

We will assess the association p-values for *rs16969968* on FEV_1_ to evaluate whether the analysis benefits from the inclusion of the regional principal components (Wang et al., 2011).

We conducted the analysis in the following way. We first prepared the genetic data from the COPDGene study for chromosome 15 with PLINK2. We employed a maximal allele frequency cutoff of *--max-maf 0.01*, LD pruning with parameters *--indep-pairwise 2000 10 0.01*, and filtered out the snp of interest using the command *--snp rs16969968*. To specify regional windows around *rs16969968*, we employed *--window W*, where we chose the window size *W* ∈ {1600, 3200, 6400}. We made sure that due to its high allele frequency (maf=0.597), the snp *rs16969968* was indeed not included in any window. Since we compute our similarity matrices on rare variants having very different allele frequencies than the common ones, they are virtually uncorrelated with the common loci that are typically tested for association in single-locus analyses. After preparing the data with PLINK2, we are left with 5,765 subjects and 94,497 RVs.

Using the genetic data above and, additionally, the covariates *age*, *sex*, an indicator of current smoking status (*smoker* =1 for current smokers and 0 for former smokers), as well as the subject’s *height* (in centimeters), we fit the regression models

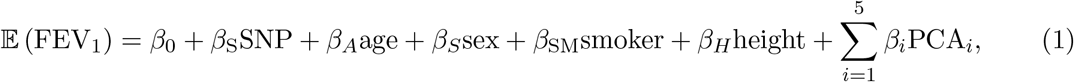

where *SNP* is the allele count data for *rs16969968*, and PCA_*i*_ for *i* ∈ {1, …, 5} are the first five principal components for either the global or regional similarity matrices. We test the hypothesis that *β*_S_ = 0 against the alternative that *β*_S_ ≠ 0. The global principal components are computed by applying any similarity matrix approach to the full genomic data, and computing the first eigenvectors. The regional principal components are computed by extracting a region around *rs16969968* (of window size given in Table 3), computing the similarity matrix on that region, and then calculating the first eigenvectors of that similarity matrix.

**Table 3:** Regression (1) on the FEV_1_ for the COPDGene study. Columns show p-values for *β*_S_ = 0 for different similarity matrices and window sizes. Five principal components each for both global and regional adjustments. Last column shows the reduction in % for the regression p-value of the combined global and regional adjustment compared to the global adjustment only.

| similarity matrix | window size | global | regional | global and regional | % reduction over global |
| --- | --- | --- | --- | --- | --- |
| Cov | 1600 | 1.81e-9 | 3.58e-9 | 1.44e-9 | 20 |
|  | 3200 | 1.81e-9 | 2.92e-9 | 1.08e-9 | 40 |
|  | 6400 | 1.81e-9 | 2.92e-9 | 1.08e-9 | 40 |
| Jaccard | 1600 | 1.82e-9 | 1.40e-9 | 9.90e-10 | 46 |
|  | 3200 | 1.82e-9 | 1.53e-9 | 1.06e-9 | 42 |
|  | 6400 | 1.82e-9 | 1.53e-9 | 1.06e-9 | 42 |
| s-matrix | 1600 | 1.29e-9 | 3.51e-9 | 8.61e-10 | 33 |
|  | 3200 | 1.29e-9 | 3.76e-9 | 1.18e-9 | 9 |
|  | 6400 | 1.29e-9 | 3.76e-9 | 1.18e-9 | 9 |
| GRM | 1600 | 8.61e-10 | 2.90e-09 | 5.33e-10 | 38 |
|  | 3200 | 8.61e-10 | 2.82e-9 | 4.82e-10 | 44 |
|  | 6400 | 8.61e-10 | 2.82e-9 | 4.82e-10 | 44 |

In this way, we regress FEV_1_ on the above covariates, where for global and regional adjustments we each use the first five principal components. For the combined global and regional adjustment, we add both the five global and the five regional eigenvectors to the model (1).

Results are given in Table 3. The table shows that for any of the four similarity matrices under investigation, and for any of the reported window sizes, the combined global and regional adjustment yields a p-value for the hypothesis *β*_S_ = 0 that is more significant (up to 46% smaller) than the global (or regional) adjustment alone. Though further investigations are necessary, these results hint at the fact that adjusting for both “global” and “regional” principal components can be beneficial in genetic association testing.

As we discussed above, in order to run the regional scans, we propose here to compute the similarity matrices based on rare variants (RVs) which have very different allele frequencies than the common variants and are therefore virtually uncorrelated with the common loci that are typically tested for association in single-locus analyses. We believe therefore that the “proximal contamination” effects (Salter-Townshend and Myers, 2019; Gazal et al., 2018, 2017; Thornton and Bermejo, 2014; Baran et al., 2012; Listgarten et al., 2012) can largely be avoided here. If RVs are to be tested for association, they should be excluded from the computation of the regional similarity matrices. Given the computational efficiency of *locStra*, this does not create a bottleneck in terms of computation time.

To assess the aforementioned proximal contamination effects (due to potential collinearity between test SNP and the estimate), we repeat the analysis above using all common snps on chromosome 1 of the EUR super population of the 1,000 Genome Project. For each common snp, we again fit the model in eq. (1). As before, this is done in three cases: for a global adjustment only (i.e., only the five global PCs), for a regional adjustment only (using 10 windows across the chromosome), and for a combined global and regional adjustment (with five global and five regional PCs). The global and regional eigenvectors were computed on the covariance matrix.

The resulting Q-Q plot is given in Figure 10. We see that the distribution of unadjusted p-values is consistent across the three cases of a global, regional, or global & regional analysis, and that the type 1 error is well controlled.

**Figure 10:**
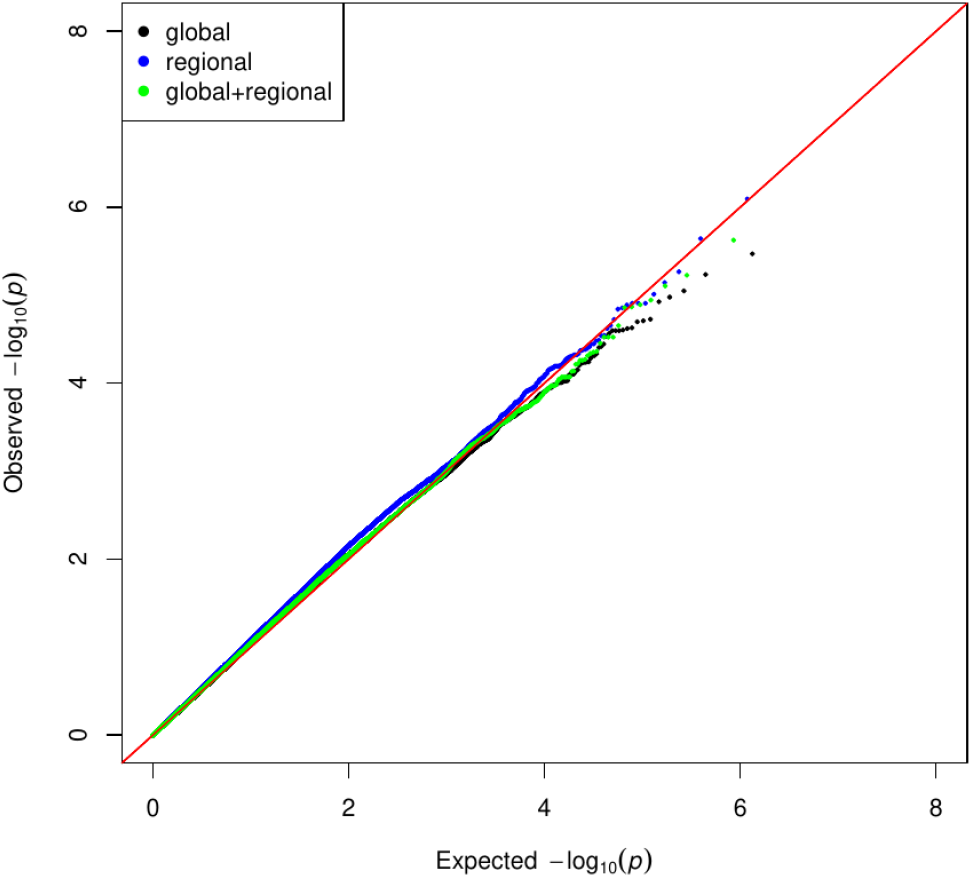
Q-Q plot for a chromosome-wide regression of all common snps using a global, regional, and global & regional adjustment according to the model of eq. (1). Chromosome 1 of the 1,000 Genome Project. Global and regional eigenvectors were computed on the covariance matrix.

Subsequent research is needed to evaluate the effects of other confounding variables, e.g. sequencing depth, batch effects, etc., on the globally and regionally computed similarity matrices. Approaches similar to the one developed for association testing (Sankararaman et al., 2008) could be utilized here.

## 4 Discussion

Our *R*-package *locStra* is the first software package that enables a comprehensive genome-wide analysis of regional stratification based on similarity matrices in WGS studies. Given a runtime of around 500 seconds for the genome-wide analysis of all sliding windows in the European super population of the 1,000 Genome Project (one Intel QuadCore i5-7200 CPU with 2.5 GHz and 8 GiB of RAM), *locStra* enables the community to conduct substantive research into regional stratification patterns at a genome-wide level in their WGS studies. At the same time, it will foster methodological work in many fields of statistical genetics, including approaches on how to incorporate regional stratification information into association studies.

## Acknowledgements

Molecular data for the Trans-Omics in Precision Medicine (TOPMed) program was supported by the National Heart, Lung and Blood Institute (NHLBI). See the TOPMed Omics Support Table (Table 1) for study specific omics support information. Core support including centralized genomic read mapping and genotype calling, along with variant quality metrics and filtering were provided by the TOPMed Informatics Research Center (3R01HL-117626-02S1; contract HHSN268201800002I). Core support including phenotype harmonization, data management, sample-identity QC, and general program coordination were provided by the TOPMed Data Coordinating Center (R01HL-120393; U01HL-120393; contract HHSN268201800001I). We gratefully acknowledge the studies and participants who provided biological samples and data for TOPMed.

Parent Study-specific Acknowledgements:

- NHLBI TOPMed: Childhood Asthma Management Program (CAMP)
- NHLBI TOPMed: Genetic Epidemiology of COPD Study (COPDGene). The COPDGene project described was supported by Award Number U01 HL089897 and Award Number U01 HL089856 from the National Heart, Lung, and Blood Institute. The content is solely the responsibility of the authors and does not necessarily represent the official views of the National Heart, Lung, and Blood Institute or the National Institutes of Health. The COPDGene project is also supported by the COPD Foundation through contributions made to an Industry Advisory Board comprised of AstraZeneca, Boehringer Ingelheim, GlaxoSmithKline, Novartis, Pfizer, Siemens and Sunovion. A full listing of COPDGene investigators can be found at: http://www.copdgene.org/directory
- NHLBI TOPMed: The Genetic Epidemiology of Asthma in Costa Rica - Asthma in Costa Rica cohort (CRA)

## Data Availability Statement

The data that support the findings of this study are openly available in “NHLBI TOPMed: The Genetic Epidemiology of Asthma in Costa Rica” at https://www.ncbi.nlm.nih.gov/projects/gap/cgi-bin/study.cgi?study_id=phs000988.v3.p1 as well as “NHLBI TOPMed: Boston Early-Onset COPD Study in the National Heart, Lung, and Blood Institute (NHLBI) Trans-Omics for Precision Medicine (TOPMed) Program” at https://www.ncbi.nlm.nih.gov/projects/gap/cgi-bin/study.cgi?study_id=phs000946.v3.p1.

## A Details on the implementation

This section briefly describes two important implementation details (for computing the covariance and Jaccard matrices) employed to enable fully sparse matrix algebra. The GRM matrix (Yang et al., 2011) and the s-matrix (Schlauch, 2016) were computed as described in their respective publications. Throughout the section, the input data *X* ∈ ℝ^*m*×*n*^ is assumed to contain (genomic) data of length *m* in each of the *n* columns, one column per individual. The parameter *m* therefore represends the number of loci included in the computation of the similarity matrix and *n* is the number of study subjects. At the end of this section, theoretical runtimes of our implementations are given.

## A.1 Covariance matrix

To compute the covariance matrix in dense algebra, let *v* ∈ ℝ^*n*^ be the column means and let *Y* ∈ ℝ^*m*×*n*^ be the matrix consisting of the rows of *X* with their mean substracted. Then

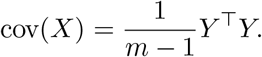

In sparse algebra, the matrix *X* cannot be normalised as in the dense case by simply substracting the column means, since this would result in a dense matrix which easily exceeds available memory. To always stay within sparse algebra, the computation is split up suitably. To be precise, let *v* denote the column means as above, and *w* ∈ ℝ^*n*^ be the column sums, then

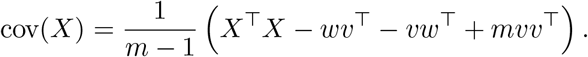

This formula has the advantage that the computation of *X^T^X* can be carried out using only one sparse matrix multiplication involving the sparse input matrix, and the remaining three outer vector products result in *n* × *n* matrices, thus never exceeding the size of the output covariance matrix.

## A.2 Jaccard similarity matrix

The entry (*i, j*) of the Jaccard matrix jac(*X*) is given as

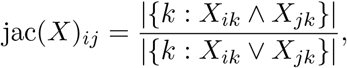

where the matrix *X* is binary.

A naïve approach to compute the entries of the Jaccard matrix loops over all entries of the Jaccard matrix and calculates the binary *and* as well as binary *or* operations on all combinations of two columns of *X*. Though having the same theoretical runtime, this naïve approach turned out to be slower in practice than the following technique which uses only one (sparse) matrix-matrix multiplication which is typically highly optimized in sparse matrix algebra packages.

Let *w* ∈ ℝ^*n*^ be the column sums of *X* as before. Compute *Y* = *X^T^X* via sparse matrix multiplication. The resulting matrix *Y* ∈ ℝ^*n*×*n*^ is dense. Compute a second matrix *Z* ∈ ℝ^*n*×*n*^ by adding *w* to all rows and all columns of −*Y*. Then, jac(*X*) = *Y*/*Z*, where the division operation is performed componentwise. This approach is computationally very fast since it relies solely on one sparse matrix multiplication, and a few more operations on the matrices *Y* and *Z* which are already of the same size as the dense Jaccard output matrix.

## A.3 Theoretical runtimes of dense and sparse implementations

Table 4 shows theoretical runtimes for both the dense and sparse matrix versions of the four similarity matrix approaches. It turns out that the runtimes for the dense computations of all similarity matrices coincide, and that the sparse computations have slightly different runtimes. The following highlights the effort of the main computation steps in each case as a function of the parameters *m* ∈ ℕ and *n* ∈ ℕ of the input data *X* ∈ ℝ^*m*×*n*^, as well as the matrix sparsity parameter *s* ∈ [0, 1] (the proportion of non-zero matrix entries).

**Table 4:** Theoretical runtimes of the four matrix approaches to compute similarity measures for both dense and sparse implementations. The runtimes are given in the parameters *m* ∈ ℕ and *n* ∈ ℕ of the input data *X* ∈ ℝ^*m*×*n*^ as well as the matrix sparsity parameter *s* ∈ [0, 1].

| method | dense | sparse |
| --- | --- | --- |
| covariance matrix | $O(mn^2)$ | $O(smn^2)$ |
| unweighted Jaccard | $O(mn^2)$ | $O(smn^2)$ |
| weighted Jaccard | $O(mn^2)$ | $O(smn^2)$ |
| GRM matrix | $O(mn^2)$ | $O(smn^2)$ |

In the dense case, computing the covariance matrix involves subtracting the column means in *O*(*mn*) and multiplying *Y ^T^Y* in *O*(*mn*^2^). In the sparse case, the computation of *X^T^X* requires *O*(*smn*^2^), and the computation of the three additional outer products requires another *O*(*n*^2^).

The Jaccard matrix involves computing *Y* = *X^T^X* in *O*(*mn*^2^) in the dense case, and *O*(*smn*^2^) in the sparse case. Adding the column sums to the rows and columns of the resulting *n* × *n* matrix takes effort *O*(*n*^2^) in both the dense and sparse case.

The effort for computing the weighted Jaccard matrix (or s-matrix) stems from the computation of weights through row sums (*O*(*mn*) in dense algebra and *O*(*smn*) in sparse algebra), multiplying the input matrix with the weights (likewise *O*(*mn*) in dense algebra and *O*(*smn*) in sparse algebra), and one matrix-matrix multiplication (*O*(*mn*^2^) in dense algebra and *O*(*smn*^2^) in sparse algebra).

Computing the GRM matrix involves the calculation of population frequencies across rows (*O*(*mn*) in dense algebra and *O*(*smn*) in sparse algebra), one matrix-matrix multiplication (*O*(*mn*^2^) in dense algebra and *O*(*smn*^2^) in sparse algebra), multiplying the input matrix with the population frequencies (*O*(*mn*) in dense algebra and *O*(*smn*) in sparse algebra), and one outer product of the population frequencies in *O*(*n*^2^).

